# The Unexpected Visibility of the SARS-CoV-2 Nucleocapsid Protein Reveals a Hidden Route of Surface Trafficking

**DOI:** 10.64898/2026.01.24.701458

**Authors:** Anthonia E. Osuagwu, Jeffrey McCausland, Christopher L. King

**Affiliations:** Department of Molecular Biology and Microbiology, Case Western Reserve University School of Medicine, Cleveland, Ohio, United States of America; Department of Pathology, Case Western Reserve University School of Medicine, Cleveland, Ohio, United States of America; Veterans Affairs Medical Center, Cleveland, Ohio, United States of America

## Abstract

The SARS-CoV-2 Nucleocapsid (N) protein, long regarded as an internal structural component of the virion, unexpectedly localizes to the plasma membrane of infected cells. Here, we show that N is actively trafficked to the cell surface via a ceramide-dependent unconventional secretory pathway. Live-stained imaging and kinetic analyses revealed that N surface association begins early in infection, before Spike (S) expression and viral release, and persists after enzymatic removal of heparan sulfate. Pharmacological disruption of phosphoinositide or phosphatidylserine interactions had minimal effect, whereas inhibition of neutral sphingomyelinase with GW4869 markedly reduced surface N, identifying a ceramide-regulated route as essential for its export. This mechanism distinguishes N from canonical transmembrane viral proteins and explains how N-specific antibodies mediate potent Fc-effector responses across SARS-CoV-2 variants. Our findings redefine the spatiotemporal dynamics of coronavirus structural proteins and reveal an unanticipated axis of immune visibility within the infected cell.

**IMPORTANCE:** Internal viral proteins are generally thought to remain confined to intracellular compartments, yet several viruses display such proteins at the surface of infected cells through mechanisms that remain incompletely defined. In this study, we characterize a host-regulated trafficking route that contributes to the delivery of the SARS-CoV-2 nucleocapsid (N) protein, a non-membrane viral protein, to the plasma membrane. Our findings indicate that N protein surface expression occurs independently of virion assembly, membrane integration, or extracellular rebinding, and instead involves host vesicular processes outside the classical secretory pathway. By elucidating how a leaderless coronavirus protein can access the cell surface, this work addresses a key gap in coronavirus cell biology and provides a framework for understanding how internal viral proteins may exploit host trafficking pathways. More broadly, these results highlight unconventional host trafficking pathways as determinants of viral protein localization and provide a framework for understanding how viruses exploit cellular export mechanisms beyond the classical secretory system.

## INTRODUCTION

For most viruses, the immune system’s recognition of infected cells is dominated by viral envelope proteins displayed on the plasma membrane. Structural proteins, typically regarded as confined to the viral interior, such as the nucleocapsid (N) of coronaviruses, are presumed invisible to extracellular immune surveillance. However, evidence from several viral systems has begun to challenge this assumption. Internal proteins from measles virus, respiratory syncytial virus (RSV), influenza virus, and retroviruses are observed at the cell surface under specific conditions, where they engage antibody-dependent effector functions. These findings suggest that immune visibility during infection extends beyond canonical membrane proteins, involving the selective externalization of intracellular antigens through unconventional trafficking mechanisms.

The N protein of SARS-CoV-2 is among the most abundant viral components in infected cells and a dominant target of T- and B-cell immunity^5–9^. Unlike Spike (S), which drives neutralization, N is an internal RNA-binding protein essential for genome encapsidation and virion assembly^10–14^. Despite lacking a signal peptide or transmembrane domain, N is detected at the cell surface of SARS-CoV-2–infected cells and in extracellular vesicle fractions, raising questions about its trafficking and function^15–18^. The immunological implications of the surface expression of N are profound: antibodies against N, though non-neutralizing, mediate potent Fc-dependent effector responses, including antibody-dependent cellular cytotoxicity (ADCC) and phagocytosis, which may contribute to cross-variant immunity. However, the cellular mechanisms by which N reaches the plasma membrane and whether its surface presence reflects active transport or passive rebinding of released protein remain unknown.

Protein secretion outside the canonical endoplasmic reticulum–Golgi pathway, termed unconventional protein secretion (UPS), provides a mechanistic framework for understanding these anomalies. Leaderless proteins exit the cell through several UPS routes: direct lipid-mediated translocation (UPS type I), ABC transporter–mediated release (UPS II), lysosomal or vesicle-based release (UPS type III), or membrane microdisruptions in stressed cells^19–21^. Many of these processes depend on specific lipids such as phosphatidylserine, phosphoinositides, or ceramide which converge at specialized membrane microdomains.

Here, we demonstrate that the N protein of SARS-CoV-2 actively traffics to the plasma membrane of infected cells via unconventional type III (UPS III) ceramide-dependent vesicles. Live-stained imaging, biochemical inhibition, and kinetic analyses show that N appears at the cell surface early in infection, before viral release, and remains detectable after enzymatic removal of glycosaminoglycans. This surface localization is sensitive to fixation but resistant to disruption of phosphoinositides or heparan sulfate, implicating dynamic lipid interactions rather than passive adsorption. Inhibition of neutral sphingomyelinase with GW4869 markedly reduces surface N, identifying ceramide-mediated vesicular trafficking as a key route. Together, our data demonstrate that N gains access to the plasma membrane, establishing an unanticipated mechanism of immune visibility for an internal coronavirus protein.

By uncovering the pathway that delivers N to the cell surface, this study reframes the immunobiology of SARS-CoV-2 infection: an internal structural protein, once thought sequestered within viral replication compartments, is repurposed as a surface antigen capable of driving Fc-effector immunity. These findings challenge long-standing assumptions about the purely structural roles of N in coronavirus assembly and open a new window into how intracellular viral components can interface directly with host immune surveillance.

## RESULTS

### SARS-CoV-2 Nucleocapsid protein is expressed at the plasma membrane with distinct distribution and kinetics from Spike protein

We previously demonstrated that SARS-CoV-2 N and S structural proteins are similarly detected at the plasma membrane surface of infected cells whereas minimal membrane (M) and no envelope (E) protein were detected^15^. To visualize the spatial distribution of these structural proteins during infection, Vero E6 cells were infected with SARS-CoV-2 at a multiplicity of infection (MOI) of 0.1 for 24 hours, and the cells were analyzed by confocal microscopy. For live-stained samples, the plasma membrane was labeled with wheat-germ agglutinin (WGA), and surface-exposed N and S were detected under non-permeabilizing conditions before fixation. After fixation and permeabilization, intracellular N was detected to confirm productive infection. Live-stained samples reveal a striking co-localization of surface N with both S and the plasma-membrane marker WGA that traces the cell surface (**Fig. 1A**). S also displays a continuous membrane staining consistent with classical ER–Golgi trafficking to the plasma membrane. This pattern indicates that N, though canonically regarded as an internal structural protein, becomes prominently exposed at the plasma membrane of infected cells. Intracellular N, detected after permeabilization, retains its diffuse cytoplasmic distribution typical of its ribonucleoprotein role. Three-dimensional z-stack reconstruction of the same live-stained cells confirmed these findings (**Supplemental Videos 1–4**). Rotating projections revealed continuous shells of S and N surrounding the cells, confirming their true surface localization rather than internal staining artifacts.

**Figure 1.**
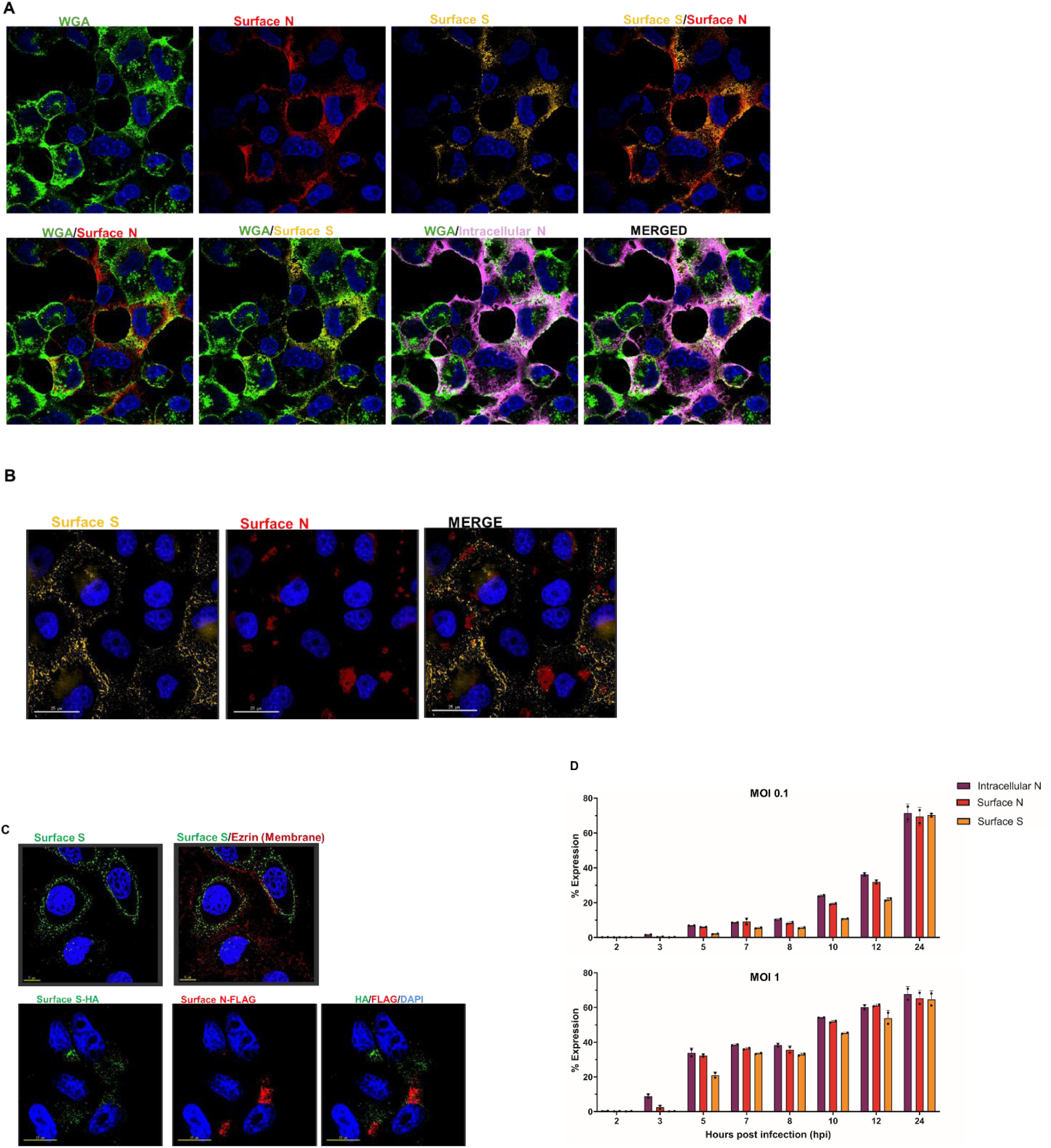
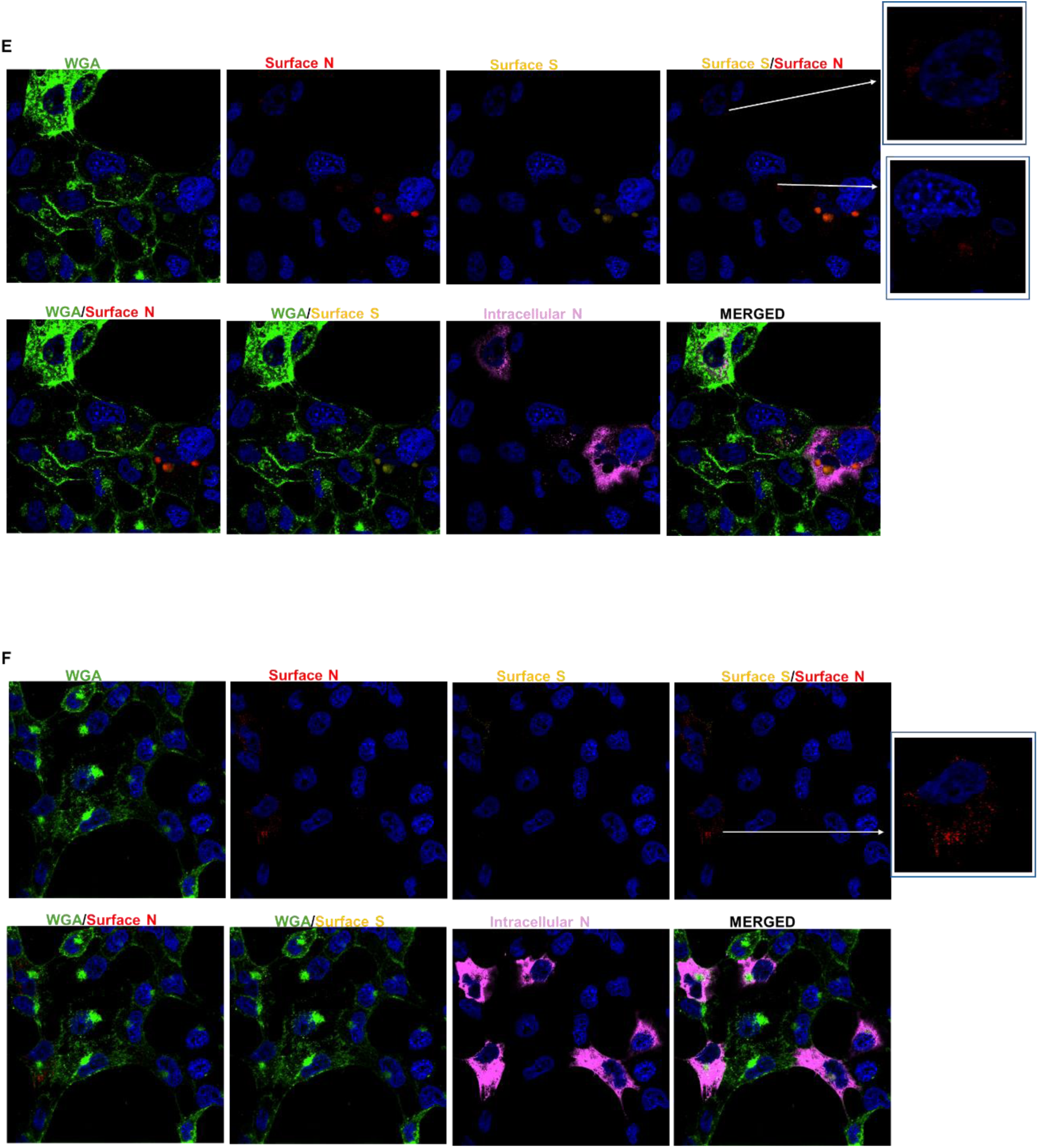
Differential distribution and kinetics of N and S at the plasma membrane. **(A)** Vero E6 cells were infected with SARS-CoV-2 (WA1) at MOI of 0.1 for 24 hrs and live-stained under non-permeabilizing conditions to detect surface N (red) and S (yellow) proteins together with the plasma membrane marker WGA (green). Nuclei were stained with DAPI (blue). After fixation and permeabilization, intracellular N (magenta) was detected. Scale bar 15μm. **(B)** SARS-CoV-2 infected Vero cells were fixed prior to antibody staining under non-permeabilizing conditions. Cells were then labeled with antibodies to N (red) and S (yellow), counterstained with DAPI (blue), and imaged. **(C)** Immunofluorescence staining under permeabilizing conditions of A549 cells stably expressing only S (top images) stained for Ezrin (red), S (green), and DAPI (blue). A549 cells co-expressing N-FLAG and S-HA (bottom images) stained for S (green), N (red), and DAPI (blue) under non-permeabilizing conditions. Scale bar 25μm. **(D)** Flow cytometric analysis of infected Vero E6 cells at MOI 0.1 or MOI 1 at the indicated timepoints showing the kinetics of intracellular N (black), surface N (gray), and surface S (light grey) expression. Data is represented from two independent experiments. Live-stained confocal images of infected Vero E6 cells at **(E)** 3 hpi and **(F)** 5 hpi showing early appearance of surface N (red, indicated by the white arrow) relative to S (yellow).

Having established surface localization of these proteins, we next assessed the stability of these membrane associations. When infected cells are fixed before antibody labeling under non-permeabilizing conditions, S maintains its uniform surface distribution, whereas N staining resolves into discrete puncta confined to the cell surface (**Fig. 1B**). The disappearance of the continuous N pattern following fixation indicates that its membrane association is fixation-sensitive and likely mediated by transient or peripheral interactions rather than stable embedding within the lipid bilayer.

To extend these observations, we generated cell lines that either stably express S or co-express N (FLAG) and S (HA) ^15^. Both proteins are detected at the plasma membrane under non-permeabilizing and permeabilizing conditions, with S co-localizing with the cortical membrane marker Ezrin and N forming distinct foci near the cell periphery (**Fig. 1C**). These findings demonstrate that N reaches the plasma membrane independently of other viral structural proteins.

### Surface Nucleocapsid protein appears earlier than Spike protein in early infection

To define the kinetics of N and S surface expression during infection, we performed a time-course analysis at low (MOI 0.1) and high (MOI 1) multiplicities of infection using flow cytometry. At high MOI, surface N is detectable as early as 3 hours post infection (hpi), whereas S is not appreciably detected until ≥5 hpi (**Fig. 1D**). By 12–24 hpi, both proteins are abundantly expressed on the surface, reaching comparable levels, consistent with progressive viral protein synthesis and particle assembly. At low MOI, surface expression of both proteins is delayed, but N consistently appears earlier than S. Importantly, intracellular staining confirmed that N protein synthesis precedes its surface appearance. Live-stained images of early time points corroborated these kinetic findings (**Fig. 1E-F**). At 3 and 5 hpi, clear surface N staining (indicated by the white arrow) with less S surface expression confirms that N is the first structural protein to reach the plasma membrane during infection. These results indicate that N is expressed earlier and reaches the cell surface earlier than S, suggesting that N utilizes an alternative trafficking pathway that precedes virion assembly or the full maturation of S via the secretory system.

These findings demonstrate that the N protein reaches the plasma membrane independently and before viral assembly. Its loss of uniformity after fixation highlights a possible vesicle or lipid-mediated mode of trafficking and membrane association distinct from the stable transmembrane integration characteristic of the S protein.

### Nucleocapsid reaches the cell surface independently of extracellular release and heparan sulfate binding

A key question arising from the detection of SARS-CoV-2 N protein at the plasma membrane is whether this localization results from an active intracellular process, or from extracellular rebinding after N has been released from lysed cells or from released virus. Recombinant N has been shown to bind strongly to glycosaminoglycans (GAGs), including heparan sulfate (HS), raising the possibility that secreted or released N could adhere to the cell surface through HS-mediated interactions^17^. If accurate, this suggests that N surface staining is a passive consequence of viral egress rather than a bona fide trafficking event.

To distinguish these possibilities, we first compared the timing of viral protein release with the appearance of surface N. Western blot analysis of culture supernatants revealed that both N and S proteins became detectable only from 7 hpi at MOI 1 (**Fig. 2A**). In contrast, surface N was consistently observed as early as 3 hpi at high MOI (**Fig. 1C**). Moreover, surface N expression preceded any detectable cytopathic effect (**Fig. 2B**), indicating that N reaches the plasma membrane before viral protein release or cell lysis occurs. These findings strongly argue against extracellular rebinding as the source of surface N in infected cells.

**Figure 2.**
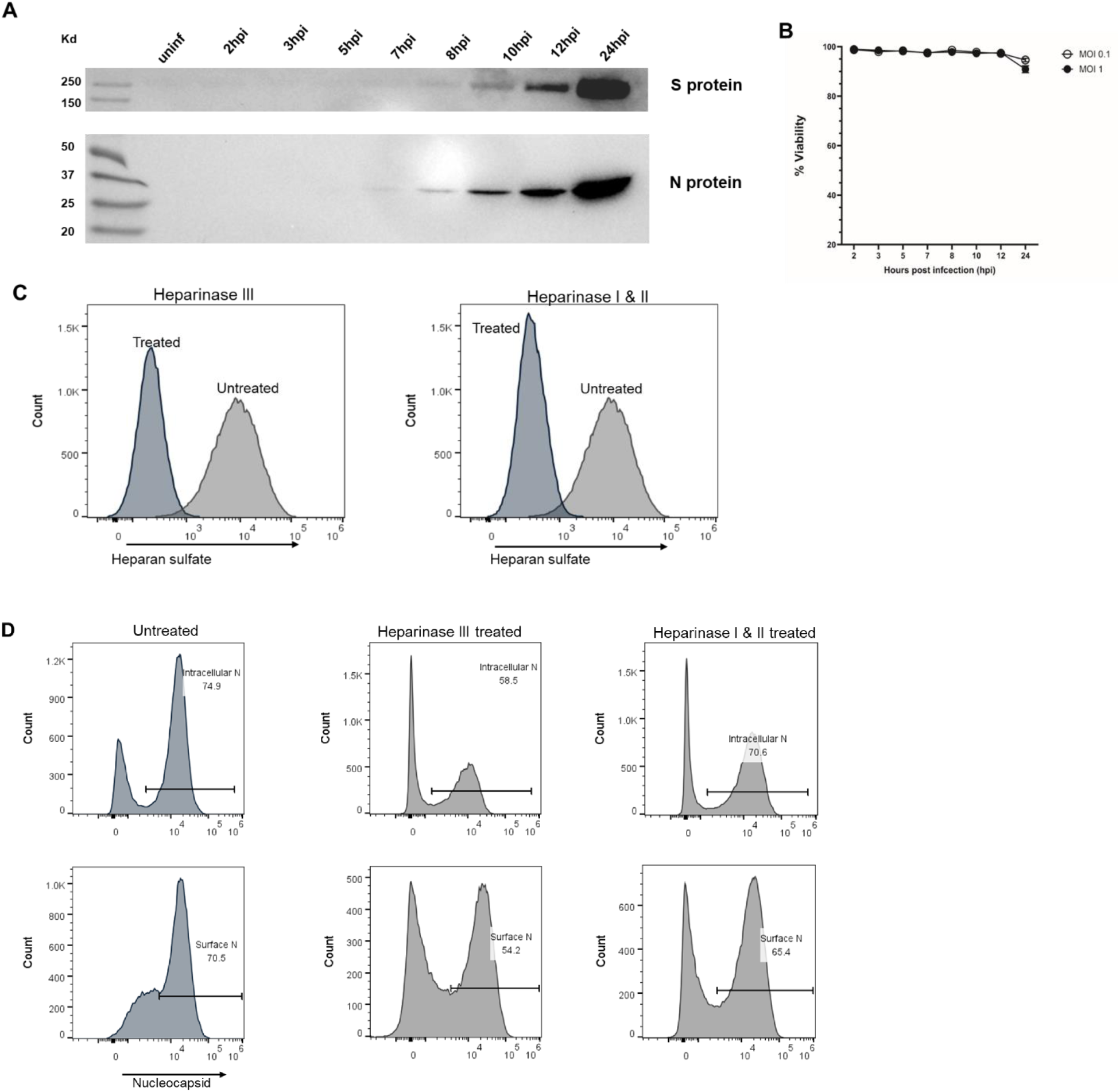
N reaches the cell surface independently of extracellular release and heparan sulfate rebinding. **(A)** Western blot analysis of SARS-CoV-2-infected Vero E6 culture supernatants collected at the indicated times post-infection. N and S proteins were detected from 7 hpi at MOI 1. **(B)** Cell viability analysis of infected Vero E6 cells measured at the indicated timepoints. **(C)** Representative flow cytometry histograms showing heparan sulfate cleavage after heparinase I, II, and III treatment of SARS-CoV-2 infected Vero E6 cells (MOI 1, 12 h post-infection). **(D)** Representative flow cytometry histograms showing intracellular N, surface N, and surface S expression in untreated or heparinase-treated infected cells (MOI 1, 12 h post-infection). The data are representative of at least two independent experiments.

We next asked whether HS maintains N at the plasma membrane. Infected cells were treated with heparinase I + II or with heparinase III, which cleaves heparan sulfate chains from proteoglycans. Enzymatic digestion effectively removed surface HS, as confirmed by flow cytometry (**Fig. 2C**). Heparinase III treatment modestly reduced the proportion of N-positive cells (intracellular N: 74.9%→58.5%; surface N: 70.5%→54.2%) (**Fig. 2C**), However, the mean fluorescence intensity (MFI) analysis showed that the surface N signal increased among remaining positive cells, while intracellular N intensity decreased (**Supplementary Table 1**). This divergence indicates that N remains robustly displayed at the plasma membrane even in the absence of HS, and the reduced frequency likely reflects secondary effects of enzymatic treatment rather than HS dependence. In contrast, treatment with heparinase I + II had no measurable effect on either the frequency or intensity of N staining (**Fig. 2D; Supplementary Table 1**).

Together, these results demonstrate that N surface exposure occurs via an active intracellular process rather than passive extracellular rebinding. Although recombinant N can bind HS in vitro, during infection, surface N appears before viral release and persists independently of heparan sulfate. These findings establish that N surface expression is a physiological feature of SARS-CoV-2-infected cells mediated by a dedicated trafficking mechanism and is not a byproduct of viral egress.

### Surface trafficking of Nucleocapsid is independent of unconventional protein secretion (UPS) type I

Having ruled out heparan sulfate rebinding as the basis for N surface detection, we next sought to determine which intracellular trafficking pathways might account for its plasma membrane localization. Recent biochemical evidence suggests that SARS-CoV-2 N interacts strongly with anionic phospholipids, including phosphoinositides and phosphatidylserine, which are enriched on the inner leaflet of the plasma membrane^22^. These interactions raise the possibility that N could exploit an unconventional protein secretion (UPS) type I mechanism, a lipid-driven route in which cytosolic proteins directly translocate across the plasma membrane through electrostatic association with membrane lipids and subsequent binding to cell-surface glycosaminoglycans (GAGs).

To test this possibility, we disrupted lipid-mediated protein trafficking with two complementary inhibitors. The first is neomycin sulfate, a polycationic aminoglycoside that binds and sequesters negatively charged phosphoinositides, thereby disrupting phosphoinositide-dependent lipid–protein interactions at the inner membrane surface. The second is polybrene, a cationic polymer that neutralizes surface anionic charges, including those contributed by glycosaminoglycans, which could interfere with electrostatic recruitment or retention of proteins at the plasma membrane.

Flow cytometric analysis revealed that neither treatment substantially altered the proportion of cells expressing surface N or S (**Fig. 3A**). Although polybrene produced a modest reduction in N-positive cells, the effect was not statistically significant, and both treatments preserved S surface expression. Live-stained immunofluorescence imaging confirms these results, showing no change in the overall distribution or intensity of surface N staining compared with untreated controls (**Fig. 3B-D**).

**Figure 3.**
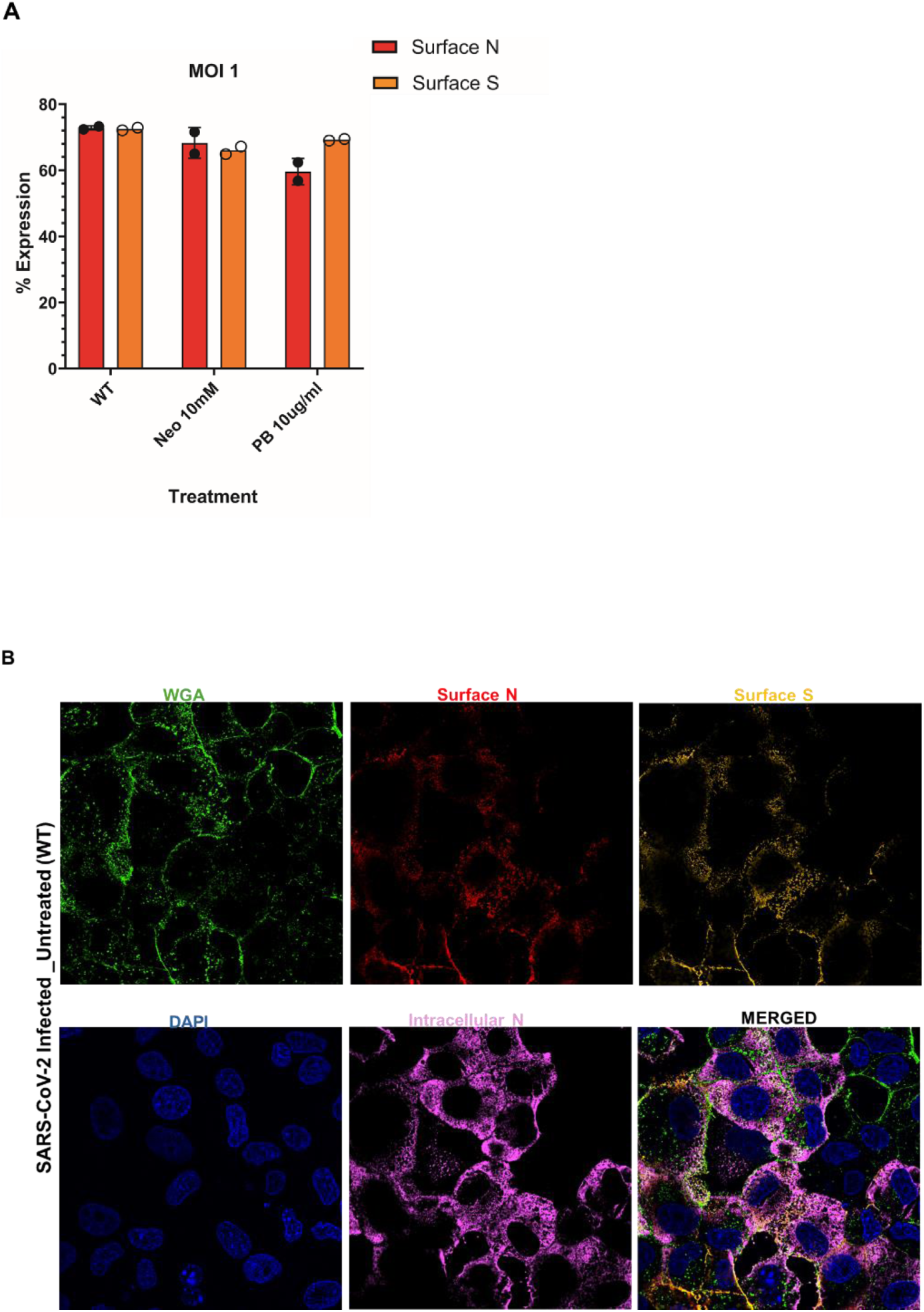
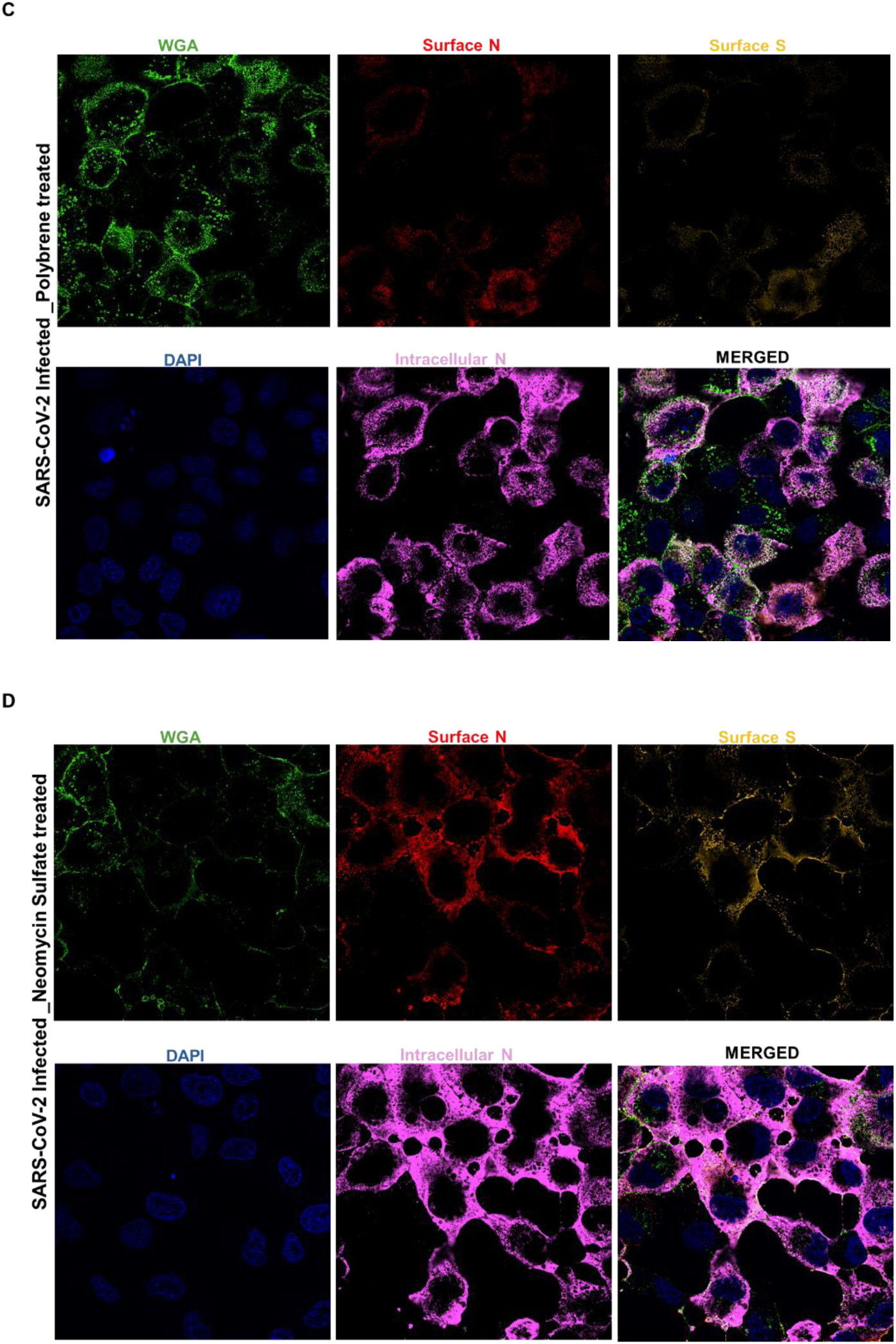
Surface trafficking of SARS-CoV-2 Nucleocapsid protein is independent of unconventional protein secretion (UPS) type I. **(A)** Flow cytometric analysis of SARS-CoV-2 infected Vero E6 cells treated with Neomycin sulfate (10 mM) or Polybrene (10 µg/mL) to disrupt lipid- and charge-mediated protein–membrane interactions. Bars show the percentage of cells expressing surface N (filled circles) and surface S (open circles). **(B–D)** Representative confocal micrographs of live-stained, infected cells showing surface N (red), S (yellow), and WGA (green) under untreated (B), Polybrene-treated (C), and Neomycin Sulfate (D) conditions. Nuclei were stained with DAPI (blue).

These findings indicate that N surface exposure does not depend on lipid-mediated translocation across the plasma membrane or electrostatic retention to GAGs on the cell surface. Instead, the data suggests that N trafficking involves a distinct intracellular route, one that bypasses the classical secretory pathway yet does not rely on the direct lipid translocation characteristic of UPS type I.

The persistence of N at the plasma membrane despite lipid-charge masking with Neomycin and Polybrene suggests that N might interact with specific lipids on the outer leaflet rather than relying on electrostatic interactions in the inner leaflet. Because phosphatidylserine (PS) is the most abundant anionic phospholipid and can transiently appear on the cell surface during membrane remodeling, we tested whether surface N co-localizes with Annexin V, which binds exposed PS. Infected cells were stained for Annexin V and N under both non-permeabilized and permeabilized conditions. In live, non-permeabilized samples, surface N appears discontinuous, with a more punctate texture than in earlier figures (**Supplemental Fig. 1**), likely because an alternative anti-N antibody recognizes a different epitope. Annexin V labeling is sparse and limited to small membrane puncta, with minimal spatial overlap with surface N, indicating that N is not detectably bound to outer-leaflet plasma membrane PS. In permeabilized cells, intracellular N is abundant in the cytoplasm, while Annexin V labels discrete cytoplasmic puncta with no co-localization. These data suggest that, although N remains membrane-associated and sensitive to fixation, its surface retention is not mediated by detectable PS binding. Instead, N may interact with other membrane lipids or protein-rich components that help stabilize its presence at the membrane surface.

### Ceramide-dependent vesicular trafficking contributes to N surface exposure

The continued presence of N surface expression despite disruption of lipid- and charge-mediated interactions suggests that its delivery to the plasma membrane occurs via an unconventional vesicle-based secretory pathway. Among such pathways, unconventional protein secretion (UPS) type III, involving lysosomal or vesicle-based release, has been implicated in the export of cytosolic proteins lacking signal peptides^23^. To test whether N exploits this vesicular mechanism, we treated SARS-CoV-2-infected cells with GW4869, an inhibitor of neutral sphingomyelinase 2 (nSMase2) that blocks ceramide synthesis and impairs exosome biogenesis, and analyzed N expression by flow cytometry and live-stained confocal imaging. Flow cytometry data show that GW4869 treatment markedly reduces the proportions of intracellular N⁺, surface N⁺, and surface S⁺ cells (Fig. 4A), consistent with broad interference with vesicular trafficking. Strikingly, only surface N lost its characteristic bimodal distribution, collapsing into a near-unimodal profile indicative of the disappearance of strongly positive cells and reduced per-cell surface intensity. In agreement, surface N MFI decreased, whereas surface S MFI and intracellular N MFI increased among the remaining positives **(Supplementary Table 2**). These divergent MFI trends suggest that GW4869 selectively impairs N export while allowing partial retention or enrichment of S at the membrane in a subset of cells.

**Figure 4.**
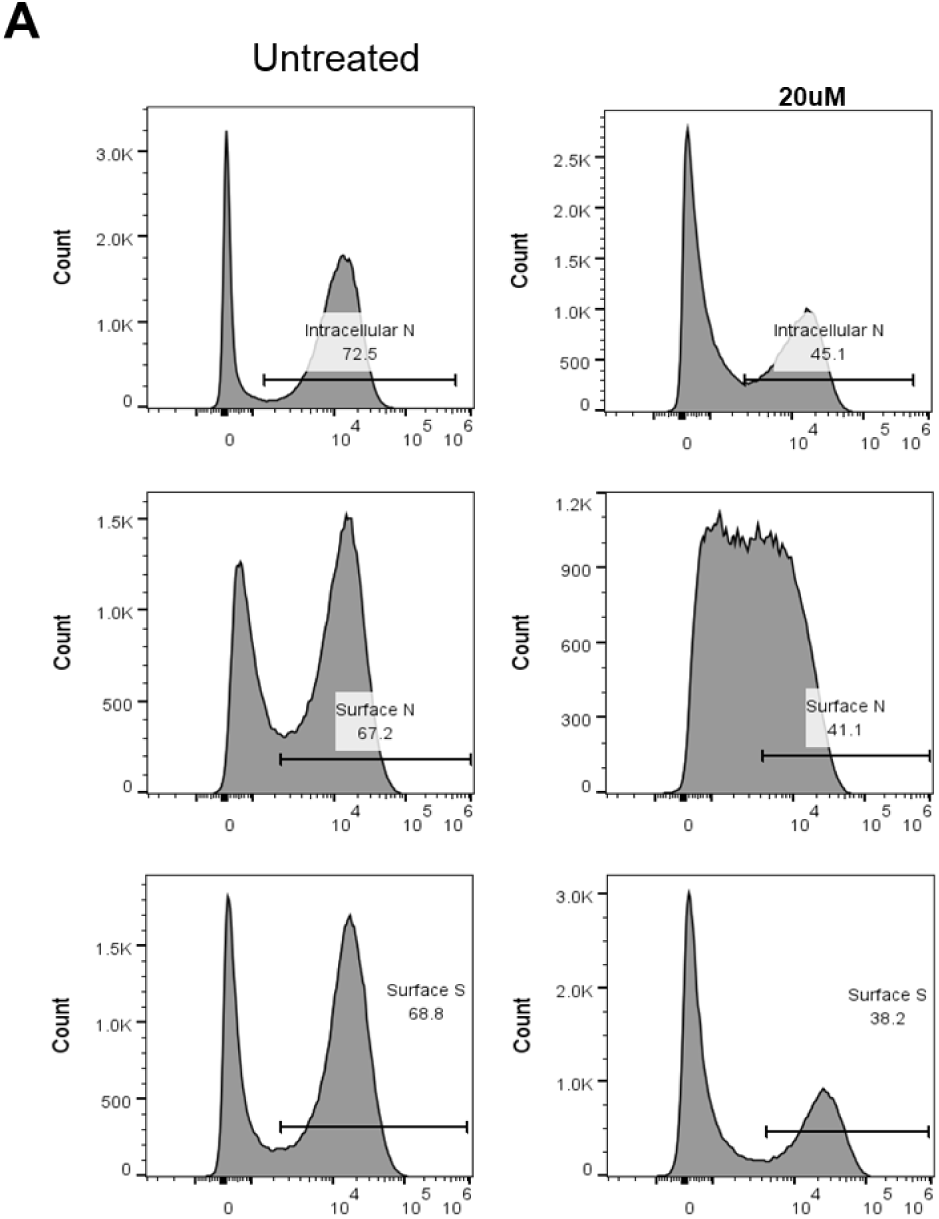
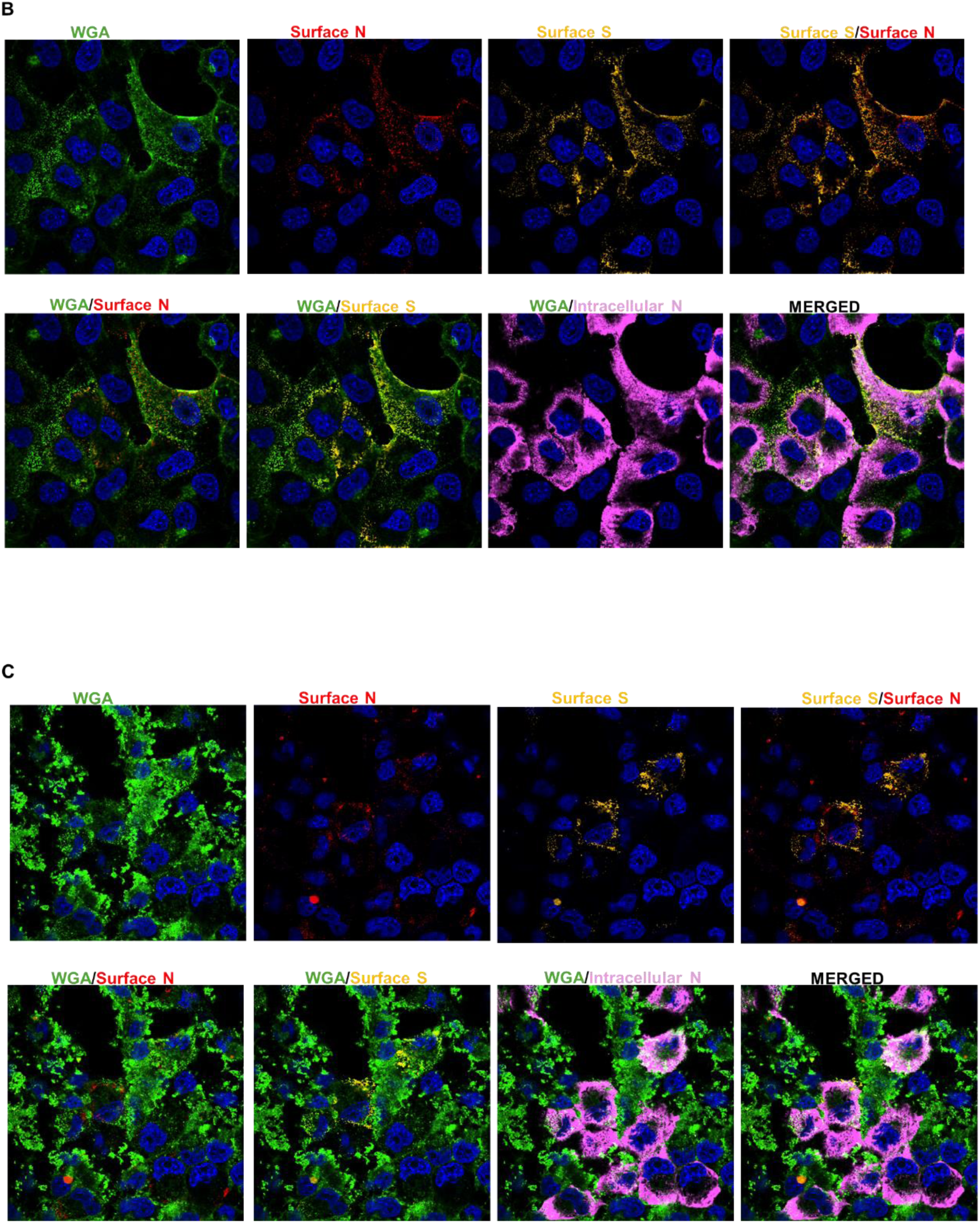
Ceramide-dependent vesicular trafficking contributes to surface N exposure. **(A)** Flow-cytometric histograms of intracellular N, surface N, and surface S in SARS-CoV-2 infected Vero cells at MOI 1, treated or untreated with GW4869 (20 µM) at 12 hpi. **(B,C)** Representative confocal micrographs of live-stained infected cells untreated **(B)** and GW4869-treated **(C)** conditions showing surface N (red), surface S (yellow), WGA (green), intracellular N (magenta), and nuclei (blue, DAPI). GW4869 reduced surface N and S signals and induced punctate WGA clusters, with surface N preferentially localizing to these regions.

Live-stained confocal imaging revealed additional insight into this interpretation (**Fig. 4B–C**). GW4869 diminished overall surface N and S staining and converted the normally continuous WGA rim into punctate, clustered microdomains. Notably, surface N but not S colocalized with these WGA-positive domains, indicating that N accumulates at ceramide or glycan-rich membrane regions that reorganize when nSMase2 activity is blocked. The reciprocal rise in intracellular N MFI supports retention of N within vesicular compartments when ceramide-dependent export is inhibited. Three-dimensional reconstructions further visualized these effects, revealing the fragmentation of WGA-positive membrane domains and the confinement of surface N within clustered glycan-rich patches upon GW4869 treatment, whereas S retained a smoother membrane distribution (**Supplementary Videos 5–8**).

Together, the loss of surface N bimodality, the decrease in surface N MFI with increased intracellular N, and the formation of clustered N-positive/WGA-positive microdomains provide converging evidence that GW4869 suppresses N trafficking to the plasma membrane, thereby blocking its surface expression. Therefore, N and S share overlapping yet mechanistically distinct trafficking pathways, with N relying on dynamic, ceramide-driven vesicles and S following the traditional secretory route.

## DISCUSSION

The present study shows that the SARS-CoV-2 N protein, long considered a purely internal structural antigen, is unexpectedly transported to and expressed on the cell surface via a regulated, non-canonical secretion pathway. This discovery adds an important mechanistic detail to the growing understanding that internal viral proteins can become recognizable to the immune system, broadening our view of viral antigen presentation beyond traditional viral envelope formation^2,24–29^. Building on our previous work showing that surface-exposed N elicits potent Fc-mediated effector responses^15^, we now define the cellular basis of this phenomenon and establish that N reaches the plasma membrane via a ceramide-dependent route independent of spike co-transport or extracellular rebinding. Together, these data identify an unanticipated trafficking pathway for a major coronavirus antigen, positioning N as both an immune target of N-specific antibodies that can trigger robust Fc-mediated effector responses against infected targets, including ADCC/ADNKA and ADCP^16,30^, and a potential participant in viral pathogenesis by sequestering chemokines^17^.

Live-stained immunofluorescence imaging showed that both N and S proteins are present at the plasma membrane of infected Vero E6 cells. N colocalizes extensively with WGA and S, indicating its exposure occurs alongside virion assembly zones or membrane microdomains. Unlike the continuous rim-like S staining, surface N appears discontinuous and sensitive to fixation, suggesting it is not stably embedded in the lipid bilayer but may associate with specific membrane lipids. This fixation sensitivity highlights a non-integral membrane interaction, likely mediated by lipid affinity or electrostatic binding rather than traditional transmembrane anchoring. The overlapping presence of N and S on the same cell surface further suggests that N exploits existing secretory or assembly interfaces, supporting the idea that viral assembly and unconventional secretion can intersect within infected cells.^31,32^.

Kinetic analyses showed that N surface exposure begins early before S surface accumulation and viral release. At high multiplicity of infection, N was detectable at 3 hours post-infection, whereas S reached similar levels only later. This timing suggests an active intracellular process rather than passive deposition or reattachment of shed viral components. Additionally, no evidence of cytopathic damage or loss of cell viability was observed during this early period, further ruling out membrane rupture as a factor. The detection of surface N before viral egress indicates that the N protein either moves directly to the plasma membrane or is temporarily exported via vesicular intermediates. This early expression likely enhances immune recognition, allowing antibody-dependent effector mechanisms even before the full formation or release of virions.

To distinguish true intracellular trafficking from extracellular rebinding, we combined kinetic analysis with heparinase treatment to evaluate the potential role of glycosaminoglycan (GAG) attachment. Although purified N protein binds strongly to heparan sulfate *in vitro*^17^, enzymatic removal of surface heparan sulfate minimally affected N surface staining and did not alter its fluorescence intensity distribution. These findings indicate that N is not secondarily adsorbed to the cell surface following release but is delivered there via a controlled intracellular route. Inhibition of the UPS type I pathway using neomycin and polybrene further demonstrated that N exposure is not caused by direct lipid flipping or by electrostatic translocation across the inner leaflet mechanisms that have been described for some leaderless secretory proteins^33–36^. Building on these observations, we then examined whether N trafficking relies on vesicular secretion driven by ceramide-rich intermediates. Mechanistically, the effect of GW4869 on both N trafficking and WGA staining can be explained by its inhibition of nSMase2, which catalyzes the hydrolysis of sphingomyelin to ceramide at the cytosolic face of the Golgi and plasma membrane. Ceramide production promotes the budding of intraluminal vesicles and the formation of exosomes and microvesicles that form the core of UPS III pathways^37,38^. By blocking this process, GW4869 not only suppresses exosome release but also perturbs the lateral organization of the plasma membrane. Ceramide depletion disrupts lipid raft integrity and alters the glycocalyx topology. WGA binds terminal N-acetylglucosamine and sialic-acid–rich glycoproteins, which are typically concentrated in these raft-like regions microdomains^39,40^. The transition from a smooth WGA rim in untreated cells to clustered WGA puncta after GW4869 treatment thus indicates fragmentation or abnormal coalescence of glycan-rich domains following the loss of ceramide-driven membrane curvature. Within these disrupted surfaces, N but not S co-localized with WGA-positive clusters, consistent with N becoming trapped at ceramide or glycan-rich patches that fail to undergo normal vesicular budding. This provides a mechanistic explanation for the observed decrease in surface N and the increase in intracellular N MFI. Therefore, inhibiting ceramide-dependent vesicle formation impedes N export while also remodeling the plasma membrane landscape where N transiently resides.

Together, these findings implicate N among the UPS type III cargoes that utilize exosome or microvesicle-like intermediates to reach the cell surface without entering the canonical ER–Golgi secretory axis^41^. Beyond this direct effect on N export, GW4869 also decreased the overall frequency of infected cells. A recent study showed that nSMase2 inhibition reduces ACE2 levels in extracellular vesicles.^42^ Because EV-associated ACE2 can enhance viral entry, its depletion likely limits infection efficiency. Thus, the profound reduction in infection we observed can be explained by a dual mechanism: impairment of N’s vesicular trafficking and loss of host EV-associated factors required for viral uptake. These observations underscore that ceramide-dependent vesicle biology contributes not only to viral protein export but also to host susceptibility, explaining the potent inhibitory effects of GW4869. Clinical studies further strengthen these findings, showing that SARS-CoV-2 proteins, including N, are packaged into circulating EVs and associate with systemic inflammation and disease severity^18,43^.

Identifying a ceramide-dependent vesicular pathway governing SARS-CoV-2 N protein surface exposure has clinical implications that extend beyond basic trafficking biology. By simultaneously limiting viral entry and suppressing the unconventional export of immunomodulatory viral proteins, host-directed inhibition of neutral sphingomyelinase–driven vesicle biogenesis could blunt early infection and downstream immune dysregulation. Although GW4869 itself is not clinically optimized, its effects highlight ceramide metabolism as a tractable target. In principle, localized modulation of this pathway, for example via intranasal or aerosol delivery to the respiratory epithelium could transiently disrupt vesicle-mediated viral processes while minimizing systemic toxicity. Such an approach would be most relevant at early stages of infection when N surface exposure precedes S presentation and may shape Fc-mediated immune engagement and chemokine sequestration. Importantly, these findings emphasize host vesicular pathways as therapeutic leverage points rather than specific drug candidates, underscoring the need for next-generation selective inhibitors that preserve epithelial integrity while limiting viral hijacking of host extracellular vesicle pathways.

These mechanistic findings align with reports across various viral families, showing that internal structural proteins can be transported to the plasma membrane via vesicular pathways. For example, the nucleoproteins of respiratory syncytial virus, measles virus, and influenza virus, as well as the Gag protein of murine leukemia virus, have been found on infected cell surfaces despite lacking signal peptides or membrane anchors ^1,44–46^. These unconventional localization events are increasingly recognized as immunologically important, providing targets for antibody-dependent cellular cytotoxicity (ADCC) and other Fc-effector mechanisms. The parallels with these systems suggest that SARS-CoV-2 N may follow a conserved evolutionary logic, leveraging host vesicular trafficking machinery to decorate the surfaces of infected cells. The fixation sensitivity of N further implies a peripheral association with lipid-enriched microdomains, potentially stabilized by phospholipids or ceramide-containing vesicles, consistent with the observed GW4869 inhibition phenotype^38^.

The immunological consequences of N surface trafficking are twofold: First, the early appearance of N at the plasma membrane may influence innate immune responses during the initial phase of infection, when viral replication and transmission occur. Because N becomes detectable at the cell surface before robust S accumulation and prior to substantial viral release, its early externalization could contribute to the induction or amplification of local innate antiviral programs. Early epithelial and innate immune activation has been shown to play a key role in limiting productive SARS-CoV-2 infection and disease progression, consistent with a model in which early antigen exposure shapes downstream immune outcomes^47^.

Second, N surface exposure fundamentally expands the antigenic landscape accessible to humoral immunity. Unlike S, which is subject to rapid mutations^48–52^, N is highly conserved and abundantly expressed^10,11,53^. Its surface presentation during infection could render infected cells recognizable to FcγR-expressing effector cells even in the absence of neutralizing antibodies. Indeed, our earlier work demonstrated that N-specific antibodies mediate robust ADCC across SARS-CoV-2 variants in the absence of S neutralizing antibodies, a property that likely reflects the conserved and early expression of N at the cell surface^15^.

In summary, our findings outline a mechanistic framework for N protein surface exposure, identifying a ceramide-dependent unconventional secretion pathway that operates independently of traditional membrane integration or extracellular rebinding. These findings challenge the conventional view of coronavirus structural protein compartmentalization and suggest that SARS-CoV-2 hijacks vesicular export pathways to display internal antigens on the cell surface. This not only increases immune visibility but may also facilitate viral spread or immune modulation through Fc engagement. More broadly, recognizing N as a dynamically trafficked surface protein broadens the understanding of coronavirus-host interactions, placing N at the interface of viral assembly, secretion, and immune detection. Future research exploring the molecular intermediaries of this pathway, including the roles of Rab proteins, ESCRT components, and ceramide-dependent microvesicle release, as well as the binding partners of surface N, will clarify how internal viral proteins reach the cell surface and redefine strategies of viral immune evasion.

### Limitations and future directions

While our study provides compelling evidence that N reaches the cell surface via a ceramide-dependent vesicular pathway, several limitations should be acknowledged. First, pharmacological inhibitors such as GW4869 and Neomycin Sulfate, although well established, are not pathway-specific and may have pleiotropic effects. Genetic approaches, such as targeted knockdown or knockout of nSMase2, anionic lipids, or other vesicle biogenesis factors, will be necessary to definitively confirm the trafficking routes involved. Second, our assays were conducted in Vero E6 cells, which offer a robust and manageable model but may not fully reflect the complexity of trafficking or immune modulation in primary human airway epithelia or in vivo. Finally, we did not directly visualize vesicle intermediates transporting N to the plasma membrane. Combining live-stained imaging with markers of multivesicular bodies or secretory autophagosomes could clarify the exact pathway.

## ACKNOWLEDGEMENTS

We thank Kien Nguyen for generously providing the cell lines expressing viral proteins used in this study. We are also grateful to Uri Mbonye for conducting earlier foundational experiments that informed the direction of this work. We thank Michael Payne for assisting with the Western blot experiment. We thank Drs. Jon Karn, Anna Bruchez, and Alan Levine for insightful scientific discussions and critical input that helped shape the experimental design and interpretation. We additionally thank Alan Levin for providing key reagents that supported this study. Finally, we appreciate the support from The Elizabeth A. Rich Biosafety Level 3 (BSL-3) Facility at Case Western Reserve University.

## AUTHOR CONTRIBUTION

A.E.O. conceptualized the study, designed and conducted the experiments, analyzed data, constructed figures, wrote and edited the manuscript. J.M. conceptualized the study, conducted initial experiments, analyzed images, and edited the original draft. C.L.K. provided intellectual and resource contributions, analyzed data, and edited the manuscript and final draft.

## DATA AVAILABILITY

All data supporting the findings of this study are included within the article and its supplementary information. Raw imaging data and quantitative analyses are available from the corresponding author upon reasonable request.

## MATERIALS AND METHODS

### Cell culture and viruses

Vero E6 cells were maintained in Dulbecco’s modified Eagle medium (DMEM; Gibco) supplemented with 10% heat-inactivated fetal bovine serum (FBS; Gibco), 1% penicillin–streptomycin (Gibco), and 2 mM L-glutamine at 37 °C with 5% CO₂. Cells were routinely tested for mycoplasma and confirmed negative. SARS-CoV-2 isolates (WT Wuhan strain) were purchased from BEI and propagated and titrated in Vero E6 cells under BSL-3 conditions using standard plaque assay protocols. Stable cell lines expressing SARS-CoV-2 viral proteins were generated using the approach previously described^15^

### Pharmacological inhibitors

Heparinase I, II (NEB), and III (Sigma) were reconstituted according to the manufacturer’s instructions and added to infected cells at 37 °C for 1 h post-infection to enzymatically remove cell-surface heparan sulfate proteoglycans. Neomycin sulfate (Sigma) was used at 10 mM, and polybrene (Sigma) at 10 µg/mL, added immediately after viral adsorption. GW4869 (Sigma; 20 µM) was dissolved in DMSO and added immediately after viral adsorption (post-treatment).

### Infection assays

Cells were seeded at 5 × 10⁵ per well in 6-well plates the day before infection. For time-course experiments, infections were performed at multiplicities of infection (MOI) 0.1 or 1.0. Virus was adsorbed for 1 h at 37 °C. The inoculum was removed and cells washed twice with PBS before incubation in fresh medium with or without inhibitors.

### Flow cytometry

At designated time points, cells were harvested with TrypLE (Gibco), washed twice in PBS, and stained with monoclonal antibodies against SARS-CoV-2 N (Sino Biological or GeneTex) and S (Sotrovimab) under non-permeabilized conditions to detect surface proteins. Cells were incubated with a fixable live-dead stain (ThermoFisher). Following fixation with 4% paraformaldehyde (PFA) for 20 min, cells were permeabilized with 0.2% Triton X-100 and stained intracellularly with anti-N. Samples were acquired on a BD LSR Fortessa or Attune NxT flow cytometer and analyzed using FlowJo v10. Percent positive cells and mean fluorescence intensity (MFI) were quantified from ≥10,000 events.

### Immunofluorescence microscopy

Cells grown on poly-L-lysine–coated coverslips were infected at MOI 0.1 or 1, washed, and live stained, then fixed or fixed after washing with 4% PFA at 12 hpi. For surface staining, cells were incubated with anti-N or anti-S (1:100 or 1:200) antibodies under non-permeabilized conditions, followed by secondary antibodies (Invitrogen, Proteintech). Wheat germ agglutinin (WGA, AF488) served as a membrane marker. For intracellular staining, cells were permeabilized with 0.2% Triton X-100 prior to antibody incubation. Nuclei were counterstained with DAPI. Images were acquired using a Nikon Eclipse Ti2 using an AXR resonant scanning detector or DeltaVision microscope using a CoolPix 1 CCD camera and processed with Nikon Elements AR software, version 6.10.02. Confocal processed images were first passed through the Nikon denoise.ai module to reduce shot noise followed by deconvolution.

### Western blotting

Cell supernatants were collected at designated times, clarified by centrifugation at 300g for 10 min, and filtered using a 0.45 µM filter (Millipore). Proteins were separated by SDS-PAGE and transferred onto PVDF membranes. Blots were probed with monoclonal antibodies against SARS-CoV-2 N (1:1,000) and S (1:1000), followed by HRP-conjugated secondary antibodies and chemiluminescent detection.

### Statistical analysis

All experiments were performed at least twice independently. Data are shown as mean ± SEM unless otherwise indicated. Flow cytometry data were analyzed using FlowJo v10 and Prism v9.

---

**Supplementary Videos 1–4** show 3D z-stack reconstructions of live-stained cells highlighting the surface localization of S (Video 1), N (Video 2), WGA (Video 3), and intracellular N (Video 4). **(Find in attached folder)**

**Supplementary Figure S1:**
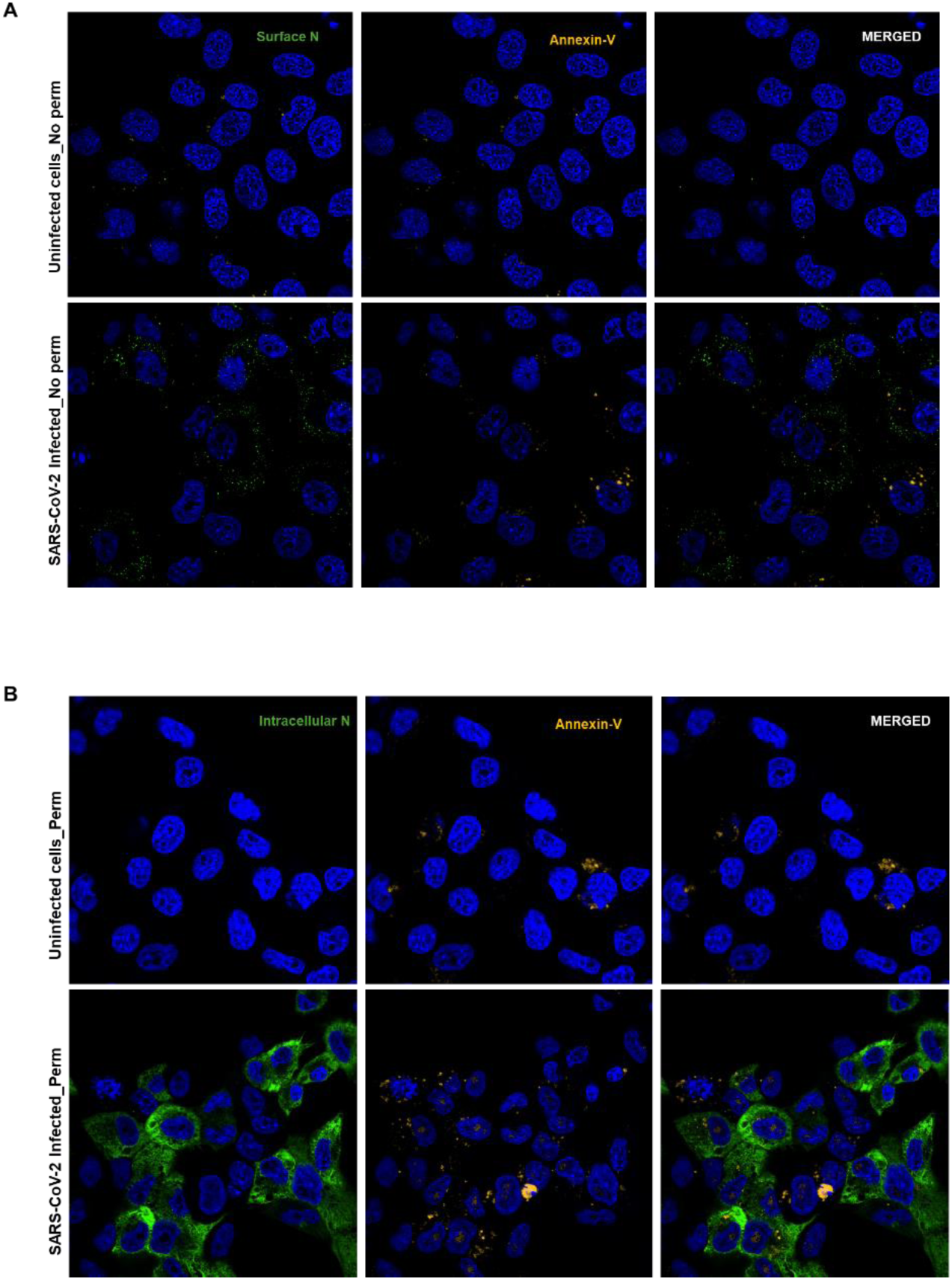
Surface N does not track with Annexin V–positive membranes. **(A)** Representative images of live, non-permeabilized staining showing surface N (green), Annexin V (yellow), and nuclei (DAPI, blue). Surface N forms a punctate rim (distinct anti-N epitope). **(B)** Permeabilized staining showing intracellular N (green) and Annexin V (yellow) in distinct puncta with no consistent co-localization.

**Supplementary Table 1.** Quantitative flow cytometry analysis of surface N, S, and intracellular N after Heparinase treatment. Quantification of the percentages of positive cells and MFI for intracellular N, surface N, and surface S in untreated and treated infected cells. Mean fluorescence intensity (MFI) among positive cells

| Samples | % Intracellular N | % Surface N | MFI Intracellular N | MFI Surface N |
| --- | --- | --- | --- | --- |
| Infected_Untreated | 74.9% | 70.50% | 16013 | 19254 |
| Infected_Hep III treated | 58.5% | 54.20% | 10039 | 24121 |
| Infected_Hep I & II treated | 70.6% | 65.40% | 16474 | 20502 |

**Supplementary Table 2.** Quantitative flow cytometry analysis of surface N, S, and intracellular N after GW4869 treatment. Infected cells were treated with GW4869 (20 µM) for 12 h prior to surface and intracellular staining and flow cytometry.

| Samples | % Intracellular N | % Surface N | % Surface S | MFI Intracellular N | MFI Surface N | MFI Surface S |
| --- | --- | --- | --- | --- | --- | --- |
| Infected_Untreated | 72.5% | 67.2% | 68.8% | 12967 | 14284 | 17638 |
| Infected_GW4869 20 uM post-treated | 45.1% | 41.1% | 38.2% | 14354 | 10959 | 25672 |

**Supplementary Videos 5–8:** Three-dimensional reconstructions of WGA-positive membrane domains and confinement of surface N within clustered glycan-rich patches upon GW4869 treatment, whereas surface S retained a smoother membrane distribution. WGA (video 5), Surface N (video 6), Surface S (video 7), Intracellular N (video 8).

